# Genome-wide association study of cerebellar volume

**DOI:** 10.1101/2021.11.04.467250

**Authors:** E.P. Tissink, S.C. de Lange, J.E. Savage, D.P. Wightman, K.M. Kelly, M. Nagel, M.P. van den Heuvel, D. Posthuma

## Abstract

Cerebellar volume is highly heritable and associated with neurodevelopmental and neurodegenerative disorders. Understanding the genetic architecture of cerebellar volume may improve our insight into these disorders. This study aims to investigate the convergence of cerebellar volume genetic associations in close detail. A genome-wide associations study for cerebellar volume was performed in a sample of 27,486 individuals from UK Biobank, resulting in 29 genome-wide significant loci and a SNP heritability of 39.82%. We pinpoint variants that have effects on amino acid sequence or cerebellar gene-expression. Additionally, 85 genome-wide significant genes were detected and tested for convergence onto biological pathways, cerebellar cell types or developmental stages. Local genetic correlations between cerebellar volume and neurodevelopmental and neurodegenerative disorders reveal shared loci with Parkinson’s disease, Alzheimer’s disease and schizophrenia. These results provide insights into the heritable mechanisms that contribute to developing a brain structure important for cognitive functioning and mental health.

## Introduction

Insights from the neuroscientific field have changed the perception that the cerebellum is predominantly involved in somatic motor control and the triad of clinical ataxias^1^. The cerebellum is now widely considered as a major centre for multiple cognitive functions such as attention^2^ and verbal working memory^3^. The association between structural abnormalities and neurodevelopmental and neurodegenerative disorders^4^ makes investigating the biology of the cerebellum an essential area of neuroscience research. Twin studies have estimated the heritability of cerebellar volume at 88%^5^. Volumetric or physical phenotypes (such as height, 73-81%^6^) often show high heritability, compared with more heterogenous phenotypes (such as psychiatric disorders, 34-79%^7^). However, it remains unclear which and specifically how genetic factors are responsible for interindividual variability in cerebellar volume.

One way to examine the genetic architecture of cerebellar volume and its overlap with neurodevelopmental and neurodegenerative disease is to evaluate common genetic variants (single nucleotide polymorphisms or SNPs) for association with neuroimaging-derived cerebellar volume variation in a genome-wide association study (GWAS). Two recent studies performing genome-wide association analysis (N = 33,224^8^ and N = 19,629^9^) on over 4,000 features of structural and functional brain organization led to the first mapping of common genetic associations, including those for the volume of cerebellar regions. The heritability based on SNPs 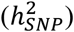 has additionally been estimated at 45.3 - 46.8%^10^, which is lower than twin-based estimates (as is also observed in other traits such as height 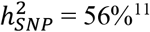 and psychiatric disorders 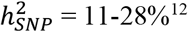). The extension of exploratory GWAS studies into the investigation of the specificity and convergence of cerebellar genetic associations is necessary to increase our understanding of the role of genetics in the cerebellum. A cerebellar volume GWAS preprint already applied functional annotation and gene mapping to identify SNPs impacting protein coding genes or gene-expression and additionally described genetic overlap between cerebellar volume and the volume of subcortical brain regions^10^. A large body of research has shown that methods that exploit the polygenic signal of traits to look for convergence onto genes, biological pathways or cell types have the potential to provide meaningful starting points for follow-up experiments^13^. In parallel, methods that zoom in on a locus level to examine the local genetic overlap between traits or pinpoint the likely causal variant(s) facilitate the prioritisation of SNPs or genes for future studies^13^. Therefore, a thorough interrogation of cerebellar volume at the level of genetic variants with detailed follow-up focussed on translating genetic loci into mechanistic hypotheses could elucidate why cerebellar volume varies between individuals and contributes to neurodevelopmental and neurodegenerative disease.

We perform a GWAS on UK Biobank data of cerebellar volume (discovery N = 27,486, replication N = 3,906) to assess which common genetic variants and genes contribute specifically to cerebellar volume. It is examined to what extent cerebellar volume-specific genes converge onto biological pathways or specific cell types located in the cerebellum and whether they display a specific temporal expression pattern through development. Further, the overlap between the genetic profile of cerebellar volume and neurodevelopmental and neurodegenerative disorders is assessed on a global and regional level. Our results provide insights into the heritable mechanisms that contribute to developing a key brain structure for cognitive functioning and mental health.

## Results

### The SNP-based heritability of cerebellar volume is 39%: 29 genomic loci identified

GWAS of cerebellar volume in 27,486 individuals assessing 9,380,224 SNPs identified 1,789 genome-wide significant (GWS) SNPs (*p* < 5×10 ^−8^) located in 29 distinct genomic loci (Figure 1a). SNP-based heritability 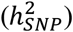 equalled 39.8 % (SE = 3.14%). The inflation of the GWAS derived test statistics (λ_GC_ = 1.18, mean χ^2^= 1.24) could be accounted for by polygenicity, since the LDSC intercept of 1.03 (SE = 0.008) indicated negligible confounding bias. We additionally calculated sample size adjusted inflation factors (λ_1,000_ = 1.006, λ_10,000_ = 1.064 and λ_100,000_ = 1.636) to allow for comparison across studies of different sample sizes. These numbers show that the expected λGC would decrease if the sample size was smaller and increase substantially if the sample size was larger, as is expected for a highly polygenic trait. 2.36% of the variance in cerebellar volume could be explained by polygenic scores (*p* = 5.86× 10^-23^) in our validation set (see Methods), using the optimal model from the target set with a *p*-value threshold of 0.29 (Supplementary Table 1). It is known that the prediction by polygenic scores can be substantially lower than the 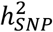 when GWAS sample sizes are low^14^, which is often the case in imaging-based GWAS. The variance explained by polygenic scores will eventually climb close to 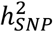 estimates if GWAS sample sizes increase, as effect sizes are then estimated with less error and differences between base and target samples will decrease^14^.

**Figure 1.**
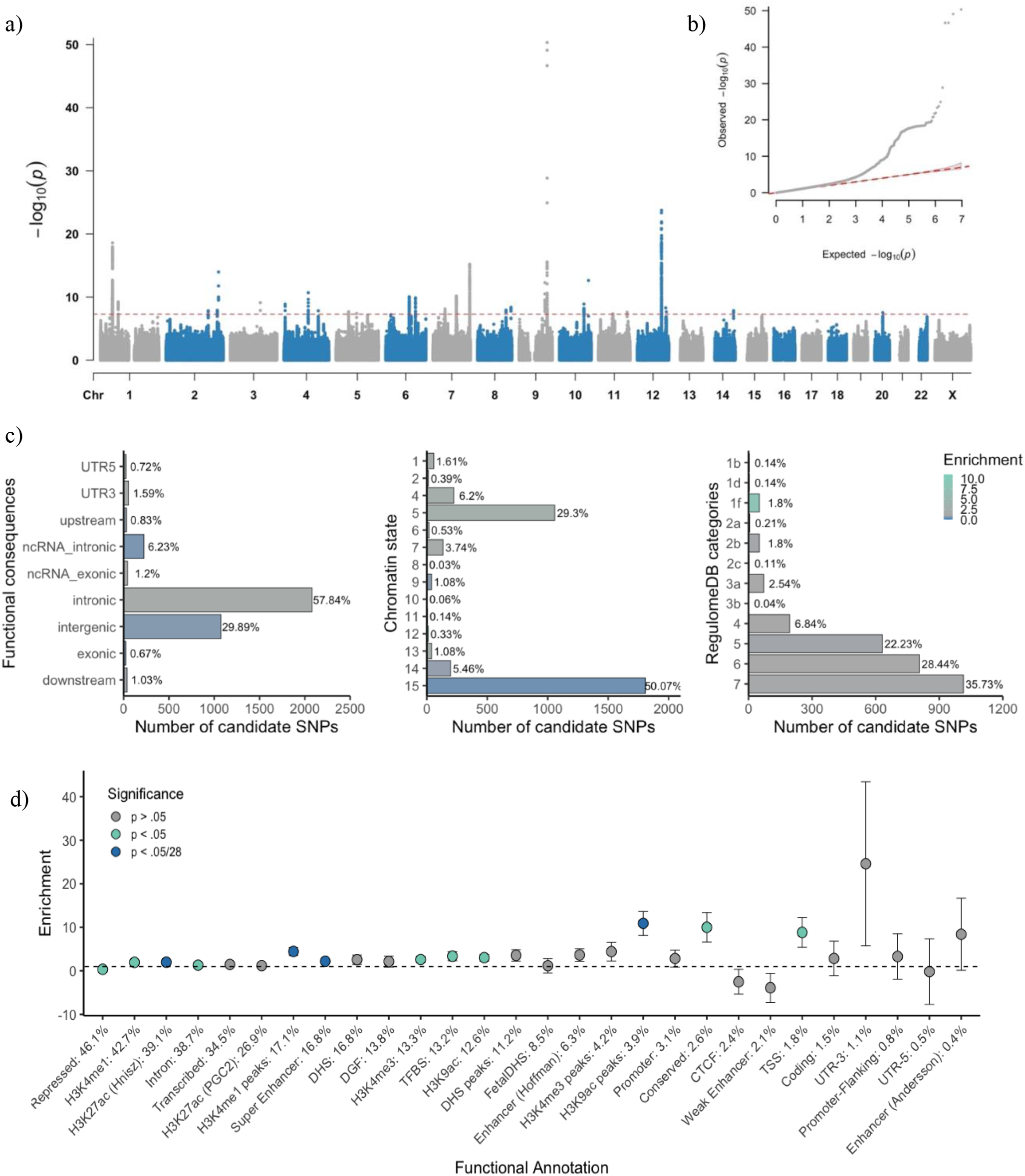
a) Manhattan plot of -log10(*p*)-values from GWAS for cerebellar volume; b) Quantile-quantile (QQ) plot of -log10(*p*)-values from GWAS of cerebellar volume; c) Proportion of candidate SNPs with corresponding RegulomeDB score, genomic location and brain-specific common chromatin state as assigned by FUMA. Enrichment values are colour-coded (depletion coloured in blue), corresponding *p*-values are available in Supplementary Table S3; d) Enrichment of cerebellar volume heritability in functional genomic categories, also available in Supplementary Table S2.

We partitioned the 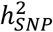 into functional genomic categories to test for enrichment of heritability in these categories (Figure 1d). With partitioned LD Score regression (LDSC) we used the associations of all SNPs, because much of the 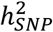 of polygenic traits lies in SNPs that do not reach genome-wide significance^15^. An enrichment of 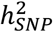 after Bonferroni correction was observed in four functional genomic categories, specifically within regions with three distinct types of chromatin modifications (Supplementary Table 2). The first category, genomic regions with H3K4me1 peaks (enrichment = 4.45, SE = 0.93, *p* = 1.45×10^-4^), is often used to distinguish enhancers from promotors^16^. In particular active enhancers are associated with additional H3K27ac^17^. Genomic regions with this H3K27ac modification (Hnisz^18^; enrichment = 1.98, SE = 0.18, *p* = 1.32×10^-7^) and super-enhancers that were tagged using H3K27ac by Hnisz *et al*^18^ (enrichment = 2.20, SE = 0.22, *p* = 4.96×10^-7^) showed significant enrichment of cerebellar volume 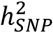 as well. Super-enhancers are large regions of regulatory elements that can affect cell type-specific and tissue-specific transcription^19^. Genomic regions with H3K9ac peaks, often active gene promoters^20^, were enriched of cerebellar volume 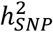 (enrichment = 10.91, SE = 2.76, *p* = 2.60× 10^-4^). Regions with these chromatin modifications show highly dynamic chromatin accessibility during both early and postmitotic stages of cerebellar granule neuronal maturation^21^. This form of chromatin plasticity is suggested to contribute to establishing synapses and, in adults, even to transcription-dependent forms of learning and memory^21^.

### Genomic location and functions of candidate cerebellar volume SNPs

We used FUMA v1.3.6a^22^ to annotate variants in associated loci based on available information about regional LD patterns and functional consequences of variants. FUMA categorized 3,611 SNPs with 1) *r*^2^ ≥ 0.6 with one of the GWS SNPs, 2) a suggestive *p*-value (< 1× 10^−5^) and 3) a minor allele frequency (MAF) > 0.005, as candidate SNPs. In concordance with our partitioned 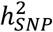 results, we observed a Bonferroni significant enrichment of candidate SNPs in brain-specific chromatin states six (genic enhancers; OR = 2.66, *p* = 1.59×10^-4^) and seven (enhancers; OR = 1.51, *p* = 9.14×10^-6^) that are both tagged by H3K4me1 modifications^23^ (for all enrichment results, see Supplementary Table 3 and Figure 1c). As is observed in other complex traits^24^, the majority of candidate SNPs were located in intronic (57.84%, OR = 1.59, *p* = 2.20×10^-149^) and intergenic regions (29.89%, OR = 0.64, *p* = 1.01×10^-92^), which complicates functional interpretation (Figure 1c). Yet four candidate SNPs were nonsynonymous exonic SNPs (ExNS; rs1060105, rs2234675, rs13107325, rs6962772) in the *SBNO1, PAX3, SLC39A8* and *ZNF789* genes respectively, of which the latter three ExNS reached genome-wide significance. *PAX3* encodes for a transcription factor that is involved in neural tube and neural crest development^25^ and is known to induce axonal growth and neosynaptogenesis in the mammalian olivocerebellar tract^26^. The missense variant in exon 6 leads to a threonine to lysine change and each effect allele accounts for a decrease in cerebellar volume of 0.17 SD. The exonic variant in the *SLC39A8* gene causes an alanine to threonine change in the zinc transporter ZIP8 the gene encodes for. Each effect allele relates to a 0.11 SD increase in cerebellar volume. The corresponding CADD scores (22.9, 25.2, 23.1, 12.6 respectively) of the four ExNS SNPs indicated high deleteriousness: a property representing reduced organismal fitness, strongly correlating with molecular functionality and pathogenicity^27^. All cerebellar volume candidate SNPs were also significantly enriched for low RegulomeDB scores (Figure 1b) (1b, OR = 15.74, *p* = 1.43e-4; 1d, OR = 17.34, *p* = 9.87e-5; 1f, OR = 18.86, *p* = 7.76e-46). Low scores represent high confidence that these SNPs have functional consequences, based on known associations with gene expression levels and the likelihood to affect transcription factor binding^28^. Supplementary Table 4 provides a detailed overview of the functional impact of all variants in the identified genomic loci.

FUMA detected 37 lead SNPs (Supplementary Table 5). We were able to replicate 20 of these 37 discovery lead SNPs in our replication sample at *p* < 0.05 and 1 of 37 at *p* < 5×10^−8^ (Supplementary Table 6). Given the winner’s curse corrected discovery associations and the sample size in the discovery and the replication phase (see Methods), we expected to find 13 (exact 13.12) and 1 (1.17) SNPs at α = 0.05 and α = 5×10^−8^, respectively. The observed replication numbers corresponding and higher to the expected level emphasize the presence of replicable genetic signal, especially at a sub-GWS level.

Although our lead SNPs represent independent variants most strongly associated with cerebellar volume, lead SNPs are not necessarily causal^29^ and could simply be correlated with the true causal SNP that was not measured on the microarray. Statistical fine-mapping can help to identify the underlying causal SNP(s), hence we applied the Bayesian fine-mapping tool FINEMAP^30^ to the 29 loci with a median number of variants of 388 per locus. The median number of variants in the 95% credible sets was 233 per locus. We used a stringent posterior inclusion probability (PIP) threshold of > 95% resulting in the selection of four credible causal SNPS (rs111891989, rs118017926, rs72754248, rs2234675) in four distinct loci. These SNPs had the greatest possibility to explain the association with cerebellar volume and overlapped with four lead SNPs identified in FUMA. The intronic variant rs72754248 in the *PAPPA* gene was the variant with the strongest association in the cerebellar volume GWAS (*p* = 4.73×10^-51^). *PAPPA* encodes for the pregnancy-associated plasma protein A, which can cleave insulin-like growth factor binding proteins (IGFBP) to regulate IGF1 availability. rs111891989 is located in an intron of *MSI1*, that encodes for an RNA-binding protein which binds to ROBO3 to regulate midline crossing of pre-cerebellar neurons^31^. rs118017926 is an intronic variant, located in the *ZNF462* gene that encodes for zinc-finger transcription factor important for embryonic neurodevelopment^32^. The fourth credible rs2234675 corresponds to the ExNS SNP in *PAX3* that was discussed above. Supplementary Table 7 includes the characteristics of all SNPs from the 95% credible sets. Figure S1 includes LocusZoom plots for the top 3 significant loci, indicating FINEMAP credible SNPs, lead SNPs and eQTLs.

### Top genes implicated in cerebellar volume suggest role for IGF1 regulation

While associated variants were mapped to 189 genes based on genomic position (located <10 kb from or within a gene (Supplementary Table 8), we also used the full GWAS results to conduct a gene-based analysis using MAGMA (Figure 2). This resulted in 85 significant genes (*p* < 0.05 / 18,852 genes tested) associated with cerebellar volume (Supplementary Table 9). 61 of these were also part of the 189 that were implicated because of the presence of a GWS SNP. The five most strongly associated genes with cerebellar volume included *PAPPA (p* = 3.37×10^-44^, confirming our SNP results discussed above), *WASHC3* (*p* = 2.50×10^-27^), *PARPBP* (*p* =2.5151e-25×10^-25^), *IGF1*(*p* = 6.85×10^-23^) and C1orf185 (*p* = 3.51×10^-22^). We also linked GWS SNPs to genes based on whether the SNP was known to be associated to cerebellum-specific expression of that gene (expression quantitative trait loci; eQTL). This resulted in the mapping of 1,197 SNPs to 32 genes for which eQTL associations in cerebellar tissue were available (Supplementary Table 8). Nineteen of these eQTL cerebellum genes overlapped with the positional strategies mentioned above, 13 were additionally discovered.

**Figure 2.**
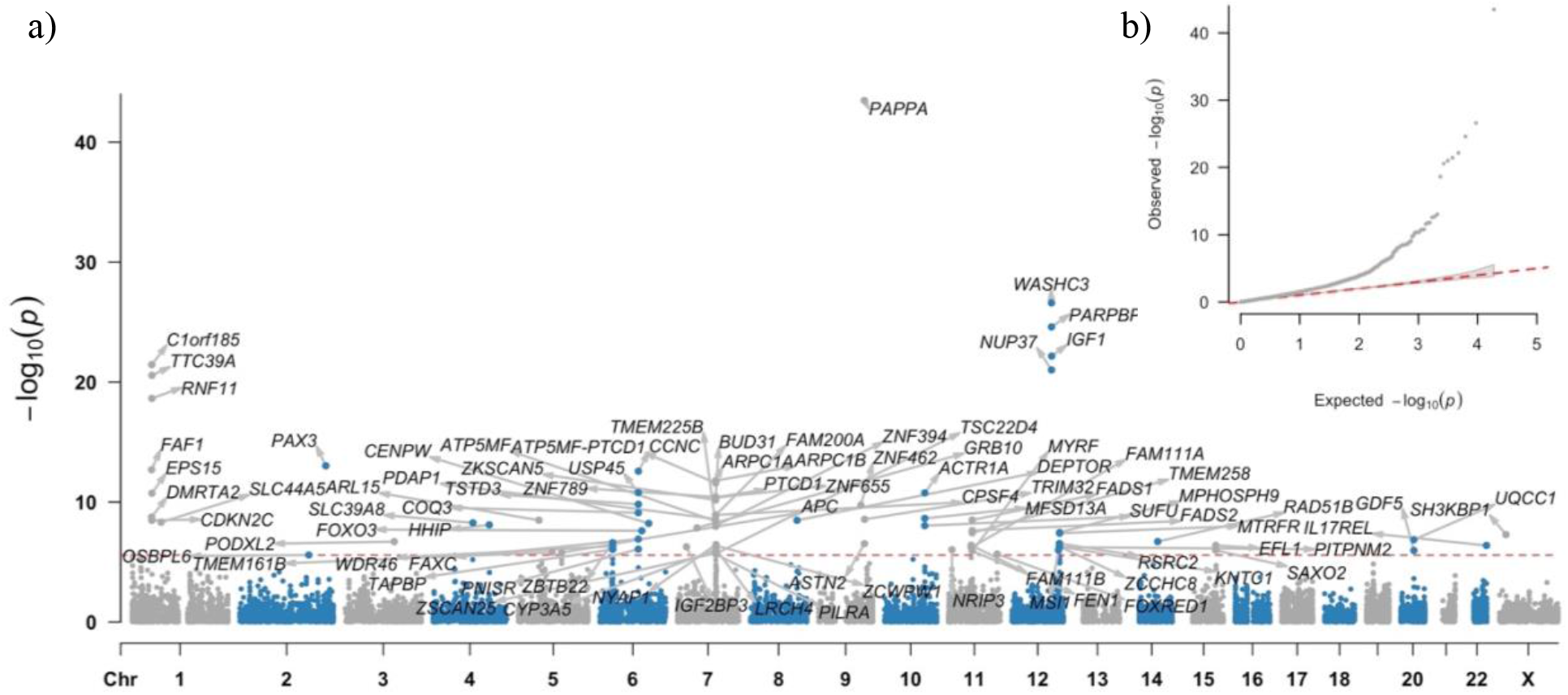
a) Manhattan plot of -log10(*p*)-values from the gene-based GWAS for cerebellar volume in MAGMA; b) Quantile-quantile (QQ) plot of -log10(*p*)-values from gene-based GWAS of cerebellar volume.

### Convergence of cerebellar volume genes on pathways, cell types and developmental stage

Next, we investigated whether associated variants and genes converged on pathways or cell types. To this end, we conducted gene-set analyses in MAGMA on the gene-based summary statistics using 7,246 MSigDB^33^ gene-sets and cell type-specificity analysis in FUMA using cerebellar cell types from the DropViz Level 1 database^34^. No evidence was found for enrichment in any of the tested MSigDB gene-sets (Supplementary Table 10). Cell type specificity analysis for nine murine cell types in cerebellar tissue resulted in a nominal significant association for enrichment of associated genes in astrocytes (β = 0.01, SE = 0.006, *p* = 0.029), yet this association did not survive the Bonferroni correction of α = (0.05/9 =) 0.056 (see Figure S2a and Supplementary Table 11).

We then tested whether associated genes showed age-specific gene expression. For this purpose, fetal, infant, adolescent and adult RNA sequencing data from cerebellar cortex tissue samples of 25 different ages from the Brainspan database^35^ were examined. MAGMA gene property analysis showed that genes specifically expressed in postconceptional week 17 (β = 0.018, SE = 0.009, *p* = 0.027) and 35 (β = 0.029, SE = 0.013, *p* = 0.015) were nominally significantly associated with cerebellar volume genes, but again this association did also not survive Bonferroni correction (Figure S2 and Supplementary Table 12).

### Global and local genetic overlap between cerebellar volume and disease

Neuroimaging studies suggest substantial evidence for phenotypic correlations between cerebellar volume and neurodevelopmental disorders^36^, such as attention deficit hyperactivity disorder (ADHD), autism spectrum disorder (ASD) and schizophrenia (SCZ), as well as neurodegenerative disorders, such as Parkinson’s disease (PD)^37^ and Alzheimer’s disease (AD)^38^. Here we tested whether global genetic correlations (*r*_g_) between cerebellar volume on the one hand and SCZ, ASD, ADHD, PD, and ALZ on the other hand were significantly different from zero using LD Score Regression (Supplementary Table 13). We did not find any statistically significant global *r*_g_ between cerebellar volume and individual disorders (Figure 3).

**Figure 3.**
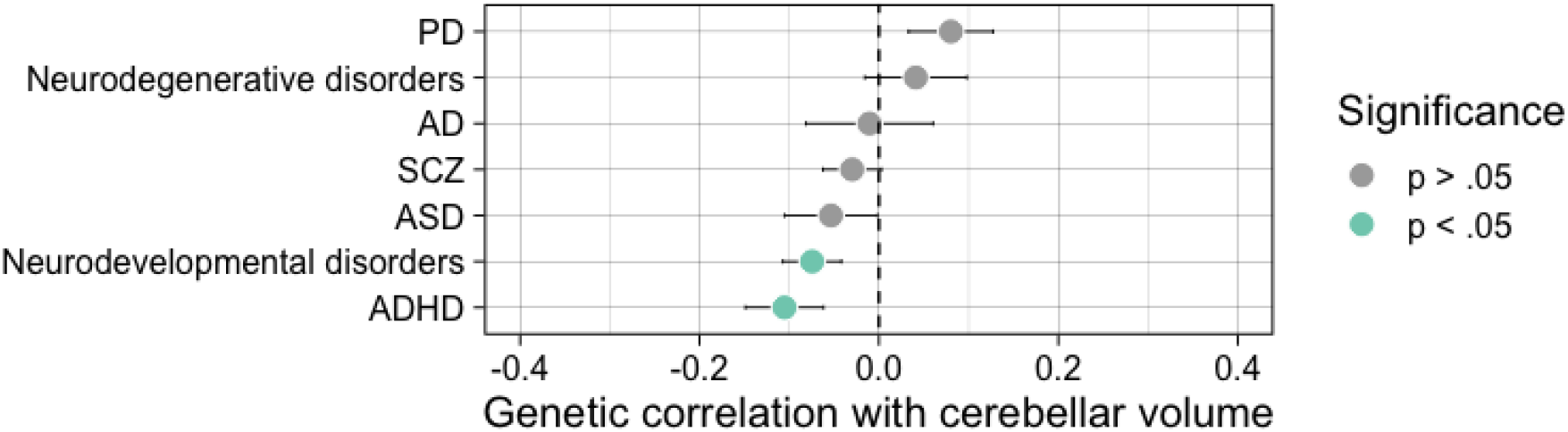
Global genetic correlations as estimated in LDSC between cerebellar volume and neurodevelopmental and neurodegenerative disorders. Although none of the traits survived Bonferroni correction for number of traits tested, we did observe a slight gradient of negative *r*_g_ with neurodevelopmental disorders to less negative and slightly positive *r*_g_ with neurodegenerative disorders. To investigate this further, we meta-analysed the ADHD, ASD and SCZ summary statistics to represent the genetic signal of the overarching neurodevelopmental disorder dimension and similarly meta-analysed PD and AD summary statistics to capture the genetic signal of the neurodegenerative disorder dimension (see Methods).

Since global *r*_g_’s are an average of local correlations across the genome, there is the possibility that substantial, but contrasting local r_g_’s are masked. To test whether this was the case, we computed local *r*_g_ in SUPERGNOVA (Figure 4 and Supplementary Table 14). Although the global *r*_g_ between cerebellar volume and PD was close to zero (*r*_g_ = 0.08, SE = 0.05, *p* = 0.089), we identified a locus on chromosome 14 that did show significant local *r*_g_ with cerebellar volume (Table 1) but did not reach GWS in both the PD and cerebellar volume GWAS. In AD and the neurodegenerative disorders meta-analysis phenotype, a positive correlation with cerebellar volume could be observed in a locus on chromosome 16 (Table 1). The lead SNP of this GWS AD locus is an intronic variant in *KAT8* and multiple significant eQTL variants in this locus influence *KAT8* expression in brain tissue including the cerebellum^39^. *KAT8* encodes for lysine acetyltransferase 8 that acetylates histone H4 at lysine 16 (H4K16ac)^40^. Alzheimer’s disease patients show large amounts of H4K16ac loss compared to normal aging, especially close to Alzheimer’s disease GWAS hits^41^. *KAT8* also seems to be vital for Purkinje cell survival. For schizophrenia we observed Bonferroni corrected significant local *r*_g_ with cerebellar volume in a positive direction on chromosome 12 and a negative direction on chromosome 19. The same loci were also Bonferroni significantly correlated between cerebellar volume and the meta-analysed neurodevelopmental disorders. The locus on chromosome 12 was GWS in both our GWAS as in the SCZ GWAS and includes 21 positionally mapped cerebellar-volume genes and 6 cerebellar-volume genes that were mapped through eQTL associations. The top hit for SCZ in this locus (rs2102949) was an intronic variant in *MPHOSPH9* that was also an eQTL in cerebellar tissue for *SETD8, CCDC62, PITPNM2* and *RP11-282O18.3. MPHOSPH9* encodes a phosphoprotein highly expressed in the cerebellum, but its function is not well understood^42^. The negatively correlated locus on chromosome 19 reached GWS in SCZ, but not in cerebellar volume. Within this locus, the most significant SCZ signal came from variants located in the *GATAD2A* gene which is considered one of the promising causal genes for schizophrenia^43^. GATAD2A is part of a protein complex named nucleosome remodelling and deacetylase (NuRD), which is an important epigenetic regulator in granule neuron synapse formation and connectivity during sensitive time windows of cerebellar development^44^.

**Figure 4.**
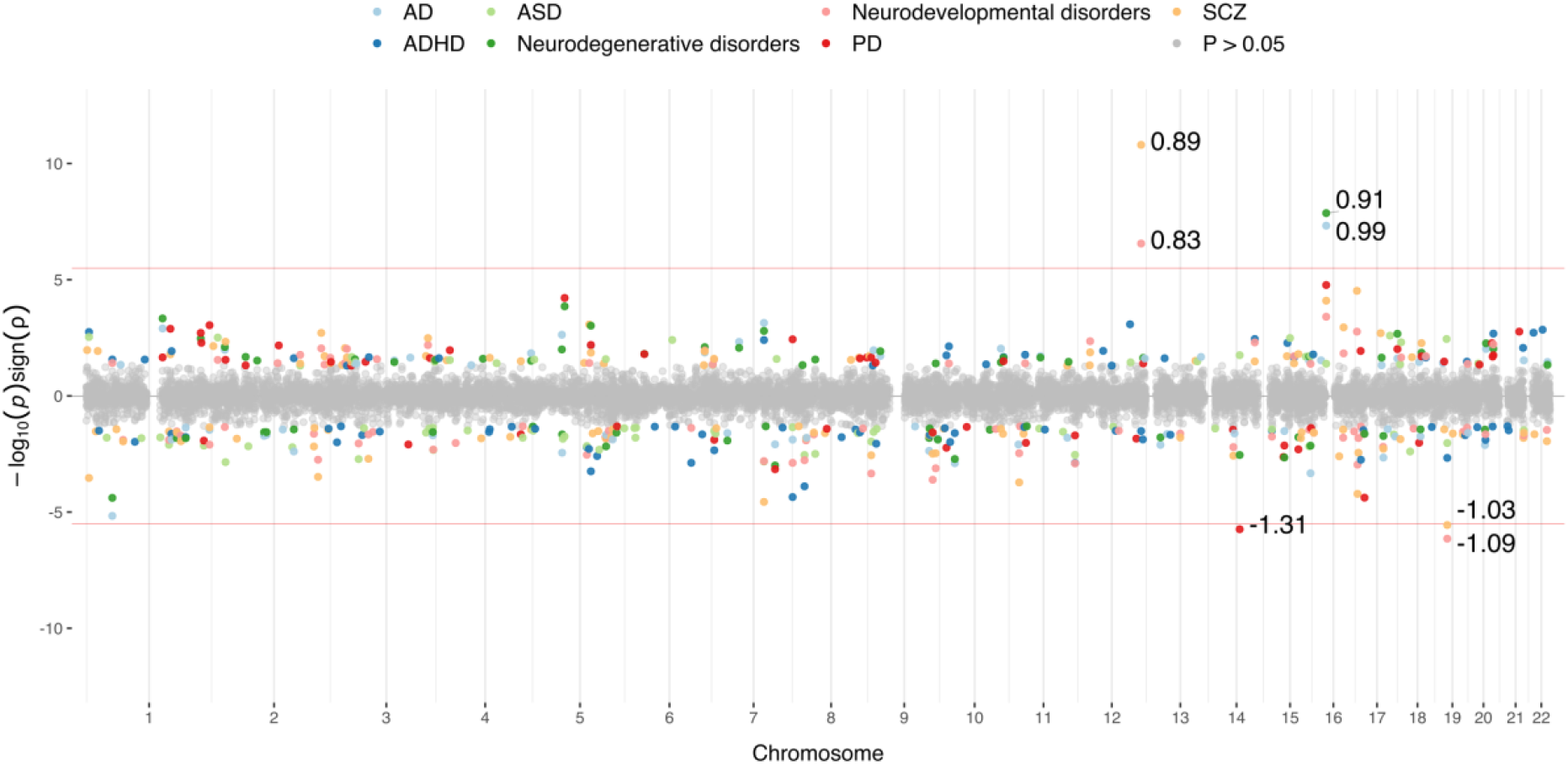
Local genetic correlations (*r*_g_) between cerebellar volume and neurodevelopmental and neurodegenerative disorders. The red line indicates the significance threshold, Bonferroni corrected for the number of loci tested in all seven traits. As during global *r*_g_, we meta-analysed the ADHD, ASD and SCZ summary statistics to represent the genetic signal of the overarching neurodevelopmental disorder dimension and similarly meta-analysed PD and AD summary statistics to capture the genetic signal of the neurodegenerative disorder dimension (see Methods).

**Table 1.** Global and local genetic correlations (rg) (Bonferroni-corrected α = 3.18×10^-6^) between cerebellar volume and PD = Parkinson’s disease, AD = Alzheimer’s disease, SCZ = schizophrenia, NDV = meta-analysis of neurodevelopmental disorders (ASD, ADHD, SCZ), NDG = meta-analysis of neurodegenerative disorders (AD,PD). Local shared SNP-heritability was calculated as 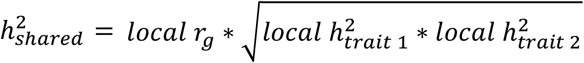.

| Trait | Locus | | | Local $r_g$ | $p$-value | Local $h_{shared}^2$ |
| --- | --- | --- | --- | --- | --- | --- |
|  | Chromosome | Start | End |  |  |  |
| PD | 14 | 67,991,028 | 69,355,606 | -1.31 | $1.83 \times 10^{-6}$ | $-3.71 \times 10^{-4}$ |
| AD | 16 | 29,694,822 | 31,380,596 | 0.98 | $4.73 \times 10^{-8}$ | $3.72 \times 10^{-4}$ |
| NDG | 16 | 29,694,822 | 31,380,596 | 0.91 | $1.36 \times 10^{-8}$ | $4.02 \times 10^{-4}$ |
| SCZ | 12 | 121,721,831 | 124,710,880 | 0.89 | $1.58 \times 10^{-11}$ | $2.43 \times 10^{-3}$ |
| | 19 | 18,506,815 | 19,873,269 | -1.03 | $2.77 \times 10^{-6}$ | $-6.81 \times 10^{-4}$ |
| NDV | 12 | 121,721,831 | 124,710,880 | 0.83 | $2.78 \times 10^{-7}$ | $1.29 \times 10^{-3}$ |
| | 19 | 18,506,815 | 19,873,269 | -1.09 | $7.16 \times 10^{-7}$ | $-6.89 \times 10^{-4}$ |

## Discussion

This study was designed to gain more insight into the genetic architecture of cerebellar volume. Our GWAS results in a substantial SNP-based heritability (39.82%) of cerebellar volume, with enrichments in super-enhancers and active promoters. The 29 GWS loci include SNPs that influence cerebellar gene expression levels or affect amino acid sequence. 85 GWS genes were found to be associated with cerebellar volume, but not specific for a cerebellar cell type, developmental stage or biological pathway. Lastly, we identify specific loci that share genetic signal between Alzheimer’s disease and Parkinson’s disease and cerebellar volume and between schizophrenia and cerebellar volume. This is the first study to stratify cerebellar volume heritability, look for convergence of the genetic signal, fine-map GWS cerebellar volume loci, and locally correlate the genetics of cerebellar volume and the neurodevelopmental and neurodegenerative disorders it is involved in.

Multiple results emphasize the importance of neurodevelopment in the genetics of cerebellar volume. The two enrichments of cerebellar volume SNPs and heritability in regions with H3K4me1, H3K27ac and H3K9ac and chromatin states six and seven (H3K4me1) stress the importance of gene regulatory enhancer elements. An animal study^21^ observed a strong overlap between regions with H3K4me1 and H3K27ac and regions with highly dynamic chromatin accessibility in cerebellar granule precursors and postmitotic cerebellar granule neurons. These dynamic regions enhance developmentally regulated increases in cerebellar granule neuron gene expression necessary for neuronal differentiation and function^21^. We highlight credible causal SNPs, exonic nonsynonymous SNPs and eQTLs in and for genes with functionalities in neurodevelopment, including *PAX3, SLC39A8, SBNO1, ZNF789, PAPPA, MSI1, ZNF462, WASHC3, PARPBP, IGF1* and *C1orf185.* Our top hit, *PAPPA*, is most strongly expressed in the placenta^45^. The last trimester of pregnancy is a critical period for cerebellum growth with a 5-fold increase in volume^46^. By cleaving insulin-like growth factor binding protein (IGFBP)-4 from IGF1, PAPPA promotes the availability of IGF1 and increases the probability that IGF1 binds its receptor^45^ and activates various intracellular signalling pathways (such as the PI3kinase-Akt pathway, promoting cell growth and maturation^47^). IGF1 receptors are most abundantly found in brain regions rich of neurons such as the olfactory bulb, dentate gyrus and cerebellar cortex^47,48^. The cerebellum hosts two-thirds of the brain’s neurons^49^. Examinations in preterm rabbit pups have indeed suggested that the loss of placental support is associated with lower levels of IGF1, decreased cerebellar external granular layer proliferation and Purkinje cell maturation^46^. Cerebellar neurogenesis also continues after birth and is accompanied with increased post-natal IGF1 levels^50^. Infant group studies have noted a positive correlation between IGF1 levels and total brain volume, with cerebellar volume showing the strongest association^51^. In mice cerebellar growth is suggested to result from increases in cell numbers^52^, as overexpression of *IGF1* caused a volumetric increase of the internal granular and molecular layer together with a respective increased number of granule and Purkinje cells^52^.

*KAT8* is another gene that influences the number of Purkinje and granular cells, as indicated by a study examining *mMof* (the mouse homolog of *KAT8*) deficient mice^53^. The current study reveals local genetic correlation between Alzheimer’s disease and cerebellar volume in a locus that includes the Alzheimer’s lead SNP in *KAT8* and multiple variants influencing *KAT8* expression in brain tissue including the cerebellum^39^. Although the study showing large amounts of H4K16ac loss in Alzheimer’s disease was performed in temporal cortex^41^, the authors note that cerebellar tissue is available for the same sample, which would be interesting to use in future research given this local genetic correlation. Genetic correlations between schizophrenia and cerebellar volume are observed in two loci that include *MPHOSPH9* and *GATAD2A* as most significant signal in schizophrenia. GATAD2A is part of the chromatin remodelling NuRD complex, which binds genome-wide active enhancers and promotors in embryonic stem cells to enable access for transcription factors to influence gene-expression^44^. Especially in the cerebellar cortex, depletion of NuRD leads to impaired development of granule neuron parallel fibers and Purkinje cell synapses^54^. Interestingly, this locus is not only associated with schizophrenia, but also with an opposing direction of effect for bipolar disorder^55^.

We also observed null results for global genetic correlations with individual disorders and for the enrichment of cerebellar volume genes in pathways, developmental stages or cell types. A possible explanation is that the substantial heritability of cerebellar volume is related to the genetic signal being divided across many genes, leading to a decreased likelihood that most of the genetic variance lies in the gene-set of interest^13^. Also note that postmortem cerebellar tissue can be scarce, for example resulting in a different single donor per developmental stage in the Brainspan database. More and more robust cerebellum specific gene-sets would therefore be interesting for future studies. Another possibility is that we did not have sufficient power to detect significant results, because of our relatively small sample size. The last decade of GWAS has proven sample size to be crucial for discovering the often small genetic effects^56^. This underscores the need for larger sample sizes for future neuro-imaging genetics studies, though this is a time consuming and costly challenge.

An important topic that needs to be addressed in neuroimaging-genetics studies is specificity. Structural properties (such as surface area, thickness, or volume) are highly correlated across brain regions, which makes finding genetic variants that go beyond global brain effects and are potentially specific to brain regions or networks challenging. In our study we included intracranial volume as a covariate in our GWAS to differentiate our findings from results on total brain volume. The authors are aware that such corrections do not guarantee pure specificity and our results reflect processes that are not exclusively involved in cerebellar volume (e.g. *IGF1* is expressed in various tissue-types and has widespread functions). However, the fact that our results can be interpreted as concordant with previous literature from the developing cerebellum or cerebellar cell types, strengthens our conclusion that these processes are involved in cerebellar volume.

A number of limitations of our study must be considered. Our sample predominantly consists of British participants of European ancestry. The effect of the European bias in the majority of GWAS remains relatively unknown, as the extent to which GWAS results can be transferred to other populations depends on many factors^57^. However, the call for a multi-ancestry GWAS approach in future studies has been gaining increasing support since it will lead to a benefit of science to all populations. Two other characteristics of our sample are the relatively high age and socioeconomic status of subjects. Genetic correlations can be influenced by genetic overlap with socio-economic status traits, as was demonstrated for mental traits previously^58^. We therefore corrected for age effects and Townsend deprivation index (TDI; a proxy of socio-economic status) within our sample, to reduce this bias in our genetic correlation analyses between cerebellar volume and neurodevelopmental and neurodegenerative disorders. Nevertheless, these sample characteristics highlight the need for other, more diverse, large neuroimaging-genetics datasets, which additionally could also aid well-powered independent replication of discovery GWAS findings. With the present data at hand, we were able to internally validate genetic variants from our discovery sample in a holdout sample to understand the generalisability of our findings.

In sum, the genetic signal for cerebellar volume measured in middle age shows a strong neurodevelopmental character. The variants and genes identified have previously been shown to influence the development of the cerebellum via chromatin accessibility of genomic elements that influence gene-expression, cell survival/death and cell growth stimulation. These insights underscore the importance of perinatal neurodevelopment for the size of a key brain structure later in life that is essential for cognitive functioning and affected in disorders.

## Online Methods

A flowchart that describes all Methods used in this manuscript is displayed in Supplementary Figure 3.

### URLs

URLs. FUMA: http://fuma.ctglab.nl/, MAGMA: https://ctg.cncr.nl/software/magma, DropViz: http://dropviz.org/, BrainSpan: http://www.brainspan.org/static/download, SynGO: https://www.syngoportal.org, winnerscurse R-package https://amandaforde.github.io/winnerscurse/, mvGWAMA: https://github.com/Kyoko-wtnb/mvGWAMA.

### Sample

All data used in this study originate from volunteer participants of the UK Biobank. The study was conducted under application number 16406. The UK Biobank obtained ethical approval from the National Research Ethics Service Committee North West–Haydock (reference 11/NW/0382) and provides researchers world-wide with a data-rich resource, including SNP-genotypes and neuroimaging data. SNP-genotypes were released for N = 488,377 participants in March 2018, with neuroimaging data available for a subsample of N = 40,682 individuals (release January 2020)^59^. In our analyses, individuals of European descent were included if their projected principal component score was closest to and < 6 SD (based on Mahalanobis distance) from the average principal component score of the European 1000 Genomes sample (n = 2,187 non-European exclusions), as has been described in previous publications by our group^60^. After stringent quality control we arrived at two randomly split samples: the discovery sample (N = 27,486) aged M = 63.55 (SD = 7.52) years with 52.50% females and the replication sample (N = 3,906) aged M = 64.91 (SD = 7.30) years with 54.58% females. Exclusion criteria were withdrawn consent, UKB-provided relatedness (subjects with the most inferred relatives, third degree or closer, were removed until no related subjects were present), discordant sex, or sex aneuploidy and resulted in n = 6,067 exclusions.

### Pre-processing and quality control

#### SNP genotype data

All participants of which data were used in this study were genotyped on the UK Biobank™ Axiom array by Affymetrix, covering 825,927 single nucleotide polymorphisms (SNPs). Both quality control and imputation were executed by the UK Biobank, using the combined Haplotype Reference Consortium and the UK10K haplotype panel as reference and resulting in 92,693,895 SNPs. Imputed variants were converted to hard call SNPs using a certainty threshold of 0.9.

Prior to downstream analyses, we performed our own additional quality control procedure. SNPs with a low imputation score (INFO<0.9), low minor allele frequency (MAF<0.005) and high missingness (>0.05) were excluded as well as multiallelic SNPs, indels, and SNPs without unique rs identifiers. This resulted in a total of 9,380,668 SNPs.

#### Neuroimaging data

T1-weighted and T2-weighted images formed the basis of the cerebellar volume estimates used in this study. The UK Biobank scanning protocol and processing pipeline is described in the UK Biobank Brain Imaging Documentation^61^. The processed and quality-controlled^62^ estimates of cerebellar white and grey matter per hemisphere were downloaded from the UK Biobank and merged to represent an estimate of total cerebellar volume. In total, n = 513 individuals had missing cerebellar volume estimates and n = 121 outliers were removed as having a cerebellar volume > 3 median absolute deviations (MAD). Mean cerebellar volume in the discovery sample was 143,580 mm^3^ (SD = 14,231 mm^3^) and 143,260 mm^3^ (SD = 14,095 mm^3^) in the replication sample, following normal distributions.

### Statistical analyses

#### Genome-wide SNP-based association study

Discovery and replication SNP-based genome-wide association studies (GWAS) were performed in PLINK version 2.00^63^ to identify and replicate common genetic variants that contribute to inter-individual cerebellar volume variability. Principal component analysis (PCA) was applied on both unrelated European neuroimaging samples using FlashPCA2^64^ to correct for population stratification. We selected a stringent set of independent (*r*^2^ < 0.1), common (MAF > 0.01), and genotyped SNPs or SNPs with very high imputation quality (INFO=1) as features in PCA (n = 145,432). The first 20 genetic principal components (PCs) served as covariates together with sex, age, genotype array, Townsend deprivation index (TDI; a proxy of socio-economic status), and several neuroimaging related confounders that were recommended by Alfaro-Almagro and colleagues^65^. These included handedness, scanning site, the use of T2 FLAIR in Freesurfer processing, intensity scaling of T1, intensity scaling of T2 FLAIR, scanner lateral (X), transverse (Y) and longitudinal (Z) brain position, Z-coordinate of the coil within the scanner, and intracranial volume (ICV). The latter is important since cerebellar volume evidently relates to the volume of the total brain, but we are interested in the genetic signal specific to the cerebellum. A linear regression model with additive allelic effects was fitted for each SNP to detect potential genetic effects on cerebellar volume. Variants on XY chromosomes were treated like autosomal variants with male X genotypes counted as 0/1 dosage. The alpha level for SNPs reaching genome-wide significance was adjusted from α=0.05 to α=(0.05/1,000,000 =) 5×10^-8^ according to the Bonferroni correction for multiple testing. Unstandardized beta values (B) were standardized (*β*) using the following equation: 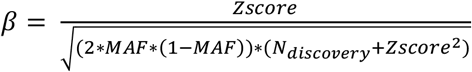, with 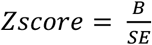.

#### (Partitioned) SNP heritability

Linkage disequilibrium score (LDSC) regression was used to estimate how much of cerebellar volume variability could be explained by additive common genetic variation^66^. This so-called SNP heritability, or 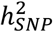, captures only the proportion of additive genetic variance due to LD between the assayed and imputed SNPs and the unknown causal variants. Precomputed LD scores based on the 1000 Genomes European data were used for this purpose.

Stratified LDSC regression was performed per functional genetic category^67^ to investigate if certain sites in the human genome contribute disproportionately to 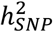 estimates. Enrichment of 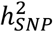 in one of the 28 categories was calculated as the proportion of 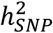 divided by the proportion of SNPs. The alpha level for significance was adjusted from α=0.05 to α= (0.05/28 =) 1.79×10^-3^ according to the Bonferroni correction for multiple testing.

#### Functional annotation and mapping of cerebellar volume associated SNPs

The summary statistics from the SNP-based GWAS served as input for the web-based platform FUMA^22^ (see URLs) to functionally map and annotate genetic associations with cerebellar volume. In order to do so, FUMA first determined which genome-wide significant SNPs were independent from one another (*r*^2^ < 0.6). SNPs in linkage disequilibrium (LD) with independent significant SNPs (*r*^2^ ≥ 0.6) were defined as candidate SNPs (using both summary statistics and 1000G Phase 3 EUR^68^). Second, a more stringent cut-off of *r*^2^ < 0.1 was used to determine which independent significant SNPs could be defined as lead SNPs. Third, genomic loci were represented by the lead SNP with the lowest *p*-value in the locus. All independent significant SNPs *r*^2^ < 0.1 with LD blocks within 250 kb distance and independent significant SNPs *r*^2^ ≥ 0.1 were assigned to the same genomic risk locus. Fourth, all candidate SNPs were annotated using ANNOVAR (1000G Phase 3 EUR as reference panel^68^), RegulomeDB score^28^ and ChromHMM^23^ (using the common chromatin states in the available adult brain samples). Enrichment of candidate SNPs falling into the categories of these annotations were calculated with Fisher’s Exact Test and *p*-values adjusted for multiple testing conformed Bonferroni correction. Fifth, annotated SNPs were mapped to one or multiple genes by two different strategies.

Positional mapping was based on physical distances (<10 kb window). Expression quantitative trait locus (eQTL) mapping was based on established associations between SNPs and a gene’s (in *cis*, < 1Mb window) expression profile in cerebellum and cerebellar hemisphere from Genotype-Tissue Expression (GTEx)^69^ v8 and cerebellar cortex from BRAINEAC^70^ databases.

#### Statistical fine-mapping of cerebellar volume associated loci

Focussing on the most significant independent SNP per locus (lead SNP, described above) is not always preferable, since the most significant variants are not necessarily causal variants^71^. Therefore, statistical fine-mapping methods have been developed to determine the probability of variants being causal. Here we applied FINEMAP^30^ to genomic loci as defined in FUMA. FINEMAP is a Bayesian statistical fine-mapping tool that estimates the posterior probability of a specific model, by combining the prior probability and the likelihood of the observed summary statistics. The posterior probabilities can in turn be used to calculate the posterior inclusion probability (PIP) of a SNP in a model and the minimum set of SNPs needed to capture the SNPs that most likely cause the association^71^. We set the maximum number of causal SNPs to 10. We additionally used LDstore^72^ to estimate the pairwise LD matrix of SNPs from quality controlled genomic data of the discovery sample. 95% credible set SNPs were defined by first summing the posterior probabilities of each model (starting from the highest probability model) until we reached a total of 95% and secondly taking all unique SNPs existing in those models. Only those SNPs with a posterior inclusion probability (PIP) > 0.95 were used for interpretation.

#### Gene-based GWAS

SNP-based GWAS approaches can suffer from power-related issues, so we combined the information from neighbouring variants within a single gene to boost power. A multi SNP-wise model for gene-based GWAS was used in MAGMA (Multi-marker Analysis of GenoMic Annotation)^73^ v1.08. The SNP-based GWAS summary statistics served as input for the gene-based GWAS and covered 18,852 genes, the UKB European population was used as an ancestry reference group. The alpha level for genes reaching genome-wide significance was adjusted from α=0.05 to α = (0.05/ 18,852 =) 2.65 ×10^-6^ according to Bonferroni correction for multiple testing.

#### Functional gene-set analysis

The summary statistics resulting from gene-based GWAS were further investigated using a pathway-based association analysis in MAGMA to test for enrichment of cerebellar volume associated genes in established biological annotated pathways. We used gene-sets from Gene Ontology (GO) molecular functions, cellular components and biological processes, and curated gene-sets from MsigDB v7.0 (sets C2 and C5)^33^. Protein-coding genes served as background genes. The alpha level for gene-sets reaching significance was adjusted from α=0.05 to α = (0.05/7,246 =) 6.90×10^-6^ according to the Bonferroni correction for multiple testing.

#### Temporal gene-set analysis

In order to identify a developmental window important for cerebellar volume we tested the relationship between age-specific gene expression profiles and cerebellar volume-gene associations (gene-based summary statistics). For this purpose, age-specific gene expression profiles were downloaded in the form of RNA sequencing data of the BrainSpan Atlas of the Developing Human Brain^35^ (URLs). We extracted gene-level RPKM values of 29 cerebellar cortex samples. From 52,376 available genes, 29,114 genes were filtered out because RPKM values were not >1 in at least one age stage, 7,828 genes were excluded because of a missing EntrezID. Finally we included genes on the SynGO^74^ list of brain genes (URLs) resulting in a set of 14,172 genes. RPKM was winsorized at 50, after which we log transformed RPKM values with pseudocount 1. Eight samples were of the same age (21 postconceptional weeks (pcw), 4 months, 3 years, 8 years), therefore log2(RPKM+1) values were averaged across samples of the same age. The log2(RPKM+1) value per gene was averaged across ages to use as a covariate. A one-sided conditional MAGMA gene-property analysis was performed, testing the positive relationship between age specificity and genetic association of genes (gene-based summary statistics).

#### Gene-set analysis in cerebellar cell types

Understanding the cell type specific expression of cerebellar volume genes may help understand the circuitry basis of cerebellar volume. We used the cell type-specificity analysis as implemented in FUMA to test whether genetic variants for cerebellar volume converge on a specific murine cerebellar cell type identified in the DropViz Level 1 database^34^. This MAGMA gene-property analysis uses gene expression values in specific cell types as gene properties and aims to test the relationship between this property and cerebellar volume-gene associations. Several technical factors, such as gene length and correlations between genes based on LD, are included to control for confounding effects^75^. The alpha level for cerebellar cell types reaching significance was adjusted from α=0.05 to α = (0.05/9 =) 5.56×10^-3^ according to the Bonferroni correction for multiple testing.

#### Global and local genetic correlations with neurodevelopment and neurodegeneration

A statistically significant genetic correlation (*r*_g_) between two traits reflects the existence of a shared genetic profile and is based on correlations in genome-wide effect sizes across traits. Cross-trait LDSC obviates the need of measuring both traits per individual, but allows for full sample overlap at the same time ^76^. Here, cross-trait LDSC regression was used to compute *r*_g_ between cerebellar volume SNP-based GWAS summary statistics and five psychiatric and neurological disorders for which well-powered (N>20,000) GWAS summary statistics were publicly available. These included the neurodevelopmental disorders autism spectrum disorder (ASD)^77^, attention deficit hyperactivity disorder (ADHD)^78^ and schizophrenia (SCZ)^79^ and the neurodegenerative disorders Parkinson’s disease (PD)^80^ and Alzheimer’s disease (AD)^39^.

Where the global *r*_g_ is an average correlation of genetic effects across the genome, local *r*_g_ can identify loci that show genetic similarity between traits. Local *r*_g_ were estimated in this study using SUPERGNOVA^81^, using the same cerebellar volume and neurodevelopmental and neurodegenerative disorder summary statistics as input. Note that, prior to local *r*_g_ estimation, SUPERGNOVA does not filter loci on univariate association signal and that local correlation estimates lower than −1 or higher than 1 may occur in loci with numerically unstable estimates of heritability^81^. The alpha level for a local *r*_g_ reaching significance was adjusted for the number of loci tested across all traits (α = 3.18×10^-6^).

After observing the results, we decided to meta-analyse the neurodegenerative disorders and meta-analyse the neurodevelopmental disorders to reduce the error term. Variants with a minor allele count (MAC) < 100 or that were not present in both (neurodegenerative disorders) or two out of three (neurodevelopmental disorders) GWAS summary statistics were excluded from the analysis. We used the software tool mvGWAMA^39^ (see URLs) to perform two effective sample size 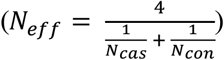 weighted analyses based on *p*-values. mvGWAMA uses the bivariate LDSC intercept to correct for sample overlap that potentially exists between the samples underlying the input GWAS summary statistics. Sample overlap between pairs of GWAS was estimated by 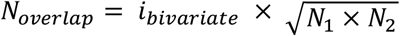 as described earlier by Yengo, Yanga & Visscher^82^, resulting in the following: ADHD/SCZ (4,457), ASD/SCZ (2,048), ADHD/ASD (18,041), AD/PD (5,438). In mvGWAMA, we performed a two-sided meta-analysis, meaning that the direction was aligned and conversion between *p*-values and z-scores was two-sided. Using the output meta-analysis summary statistics, global and local genetic correlations were estimated as described above. The alpha level for global and local genetic correlations were additionally adjusted for the number of traits tested according to the Bonferroni correction for multiple testing. For all global and local genetic correlations, the shared heritability between cerebellar volume and the other trait was calculated as 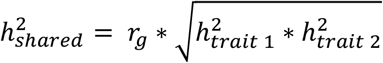.

#### Replication of discovery lead SNPs

The replicability of our GWAS results were evaluated by assessing the associations of 37 discovery lead SNPs in our replication GWAS. For this purpose, we compared observed numbers of replicated lead SNPs with numbers expected under a natural alternative hypothesis previously described by Okbay et al^83^ (Supplementary Information 1.8). Under this hypothesis, the probability that a discovery lead SNP *i* is significant in the replication sample is

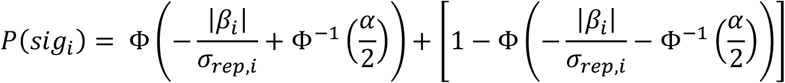

with α representing an alpha level of 0.05 and 5×10^-8^, Φ the cumulative normal distribution function, Φ^−1^ the inverse normal distribution function, *σ_rep, i_* the standard error of SNP *i* in the replication GWAS and *β_i_* the winner’s curse adjusted association estimate of SNP *i*. Winner’s curse is the occurrence of overestimated effect sizes that are induced by significance thresholding^84^. Correction for winner’s curse was performed using the Conditional Likelihood method^85^ in the winnerscurse R package (see URLs), which uses both the discovery and replication betas and standard errors for correction. Finally, the number of SNPs that is expected to show significance was defined as ∑*_i_ P*(*sig_i_*).

#### Polygenic scores

In order to estimate how much variance in cerebellar volume could be explained by our GWAS findings, we calculated polygenic scores using the discovery summary statistics (base set) in PRSice-2^86^. For this purpose, we randomly split the replication sample into a target set (N = 1,971) and a validation set (N = 1,983). A three-phase procedure was applied, with phase 1 representing the estimation of effect sizes in our discovery GWAS. In phase 2, SNP data (MAF > 0.1, chromosome X excluded) of the target sample was clumped and high-resolution *p*-value thresholding was applied using default settings to find the best fit model. All models were corrected for the same covariates as during GWAS analysis described above. During phase 3, the best fit model was fit on clumped SNP data of the validation set to be able to report the phenotypic variance explained and *p*-value unaffected by overfitting.

## Acknowledgements

D.P. was funded by The Netherlands Organization for Scientific Research (NWO VICI 453-14-005), NWO Gravitation: BRAINSCAPES: A Roadmap from Neurogenetics to Neurobiology (Grant No. 024.004.012), and a European Research Council advanced grant (Grant No, ERC-2018-AdG GWAS2FUNC 834057). The work of S.L. was supported by ZonMw Open Competition, project REMOVE 09120011910032. D.W. was funded by NWO Gravitation: BRAINSCAPES: A Roadmap from Neurogenetics to Neurobiology (grant no. 024.004.012). The work of M.H. was supported by a VIDI (452-16-015) grant from the Netherlands Organization for Scientific Research (NWO) and an ERC Consolidator of the European Research Council (101001062). The research has been conducted using the UK Biobank Resource (application no. 16406). Analyses were carried out on the Genetic Cluster Computer hosted by the Dutch National computing and Networking Services SURFsara. We thank J.P. Beauchamp for his clarifying emails about the holistic replication method.

## Author contributions

E.T., M.H. and D.P. contributed to the conception and design of the study. J.S. performed preprocessing of genetic data. E.T. performed all analyses. D.W. and K.K provided data and analysis scripts in two analyses. E.T., S.L., J.S., D.W., K.K, M.N., M.H. and D.P. contributed to the interpretation of the data. E.T. has drafted the work. E.T., M.N., M.H. and D.P. substantively revised it and J.S., S.L., D.W. and K.K. revised it.

## Data availability

Genome-wide summary statistics will be made publicly available via https://ctg.cncr.nl/software/summary_statistics/ upon publication.

---

**Supplementary Figure 1.**
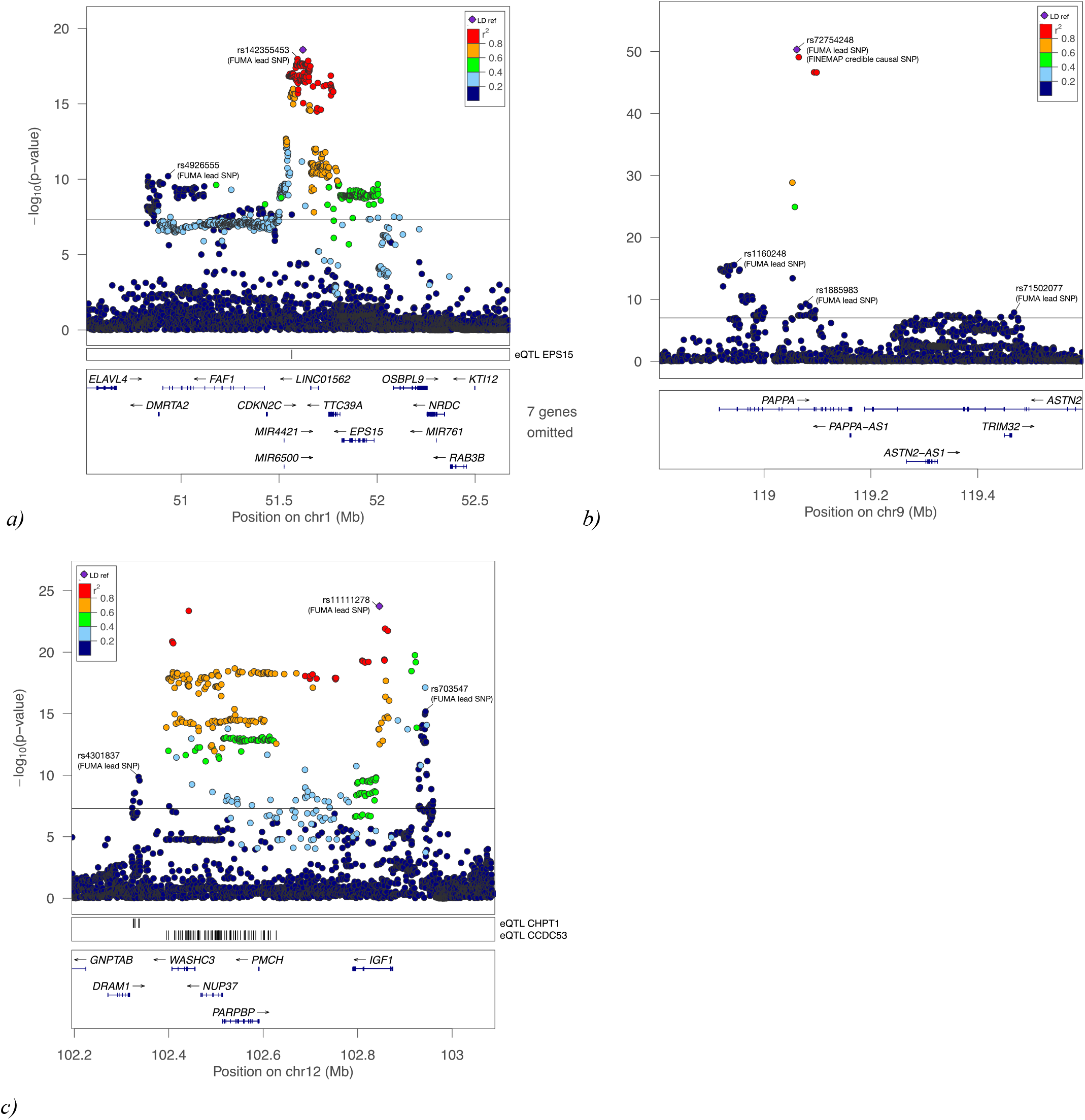
LocusZoom plots for the three most significant loci in our cerebellar-volume GWAS on chromosome 1 (a), 9 (b) and 12 (c). FUMA lead SNPs, FINEMAP credible causal SNPs (PIP > 0.95) and/or eQTLs for cerebellar tissue are indicated if applicable.

**Supplementary Figure 2.**
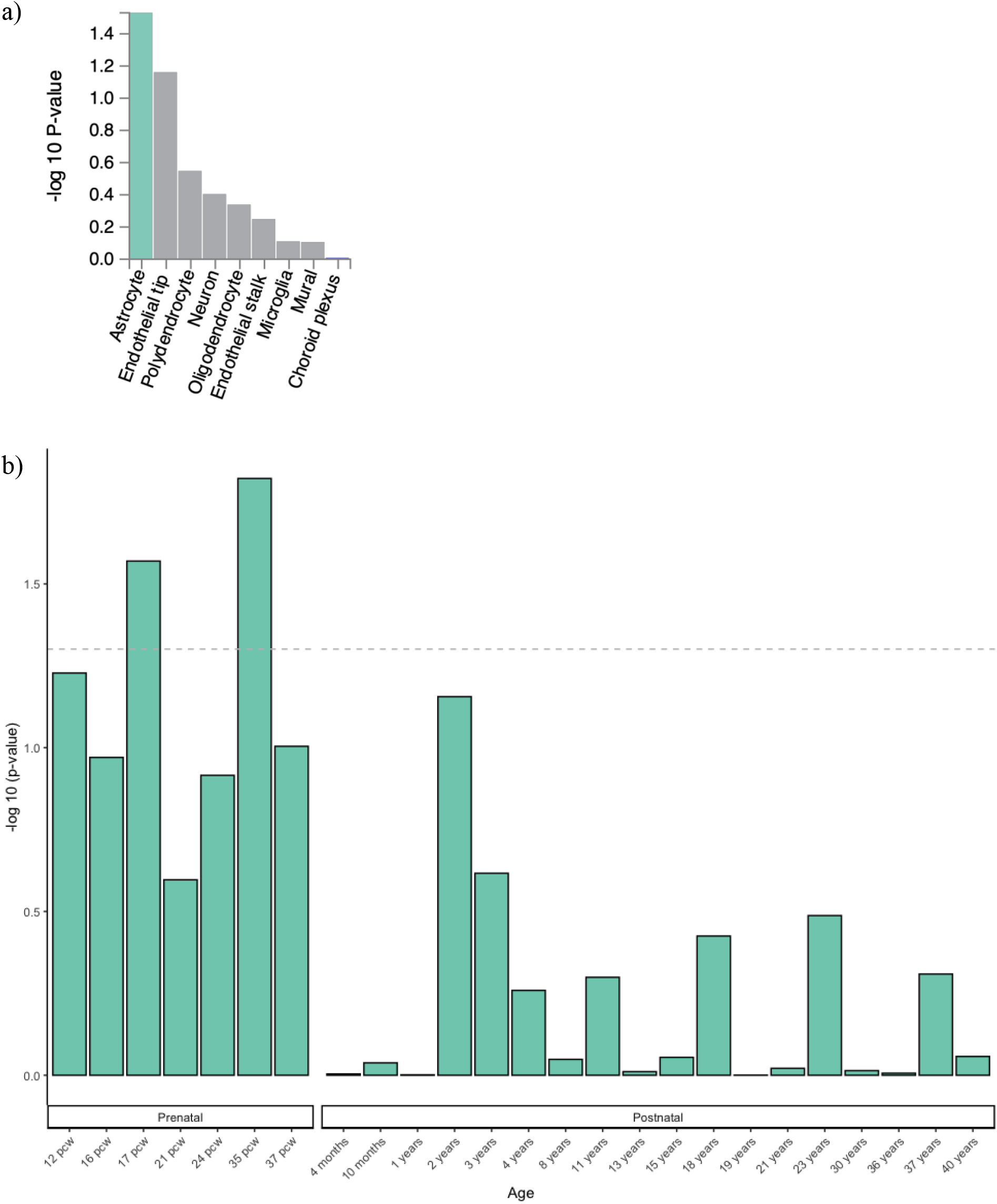
-log10(*p*)-value results from two gene-set analysis approaches: a) Gene-expression in cell types from cerebellar mouse tissue (DropViz database in FUMA) was nominally associated with the cerebellar volume gene-based GWAS sumstats in astrocytes, but this did not survive Bonferroni correction. b) Cerebellar gene-expression in donors from different developmental stages (Brainspan database) was associated nominally associated with the cerebellar volume gene-based GWAS sumstats at 17 and 35 postconceptual weeks, but these did not survive Bonferroni correction.

**Supplementary Figure 3.**
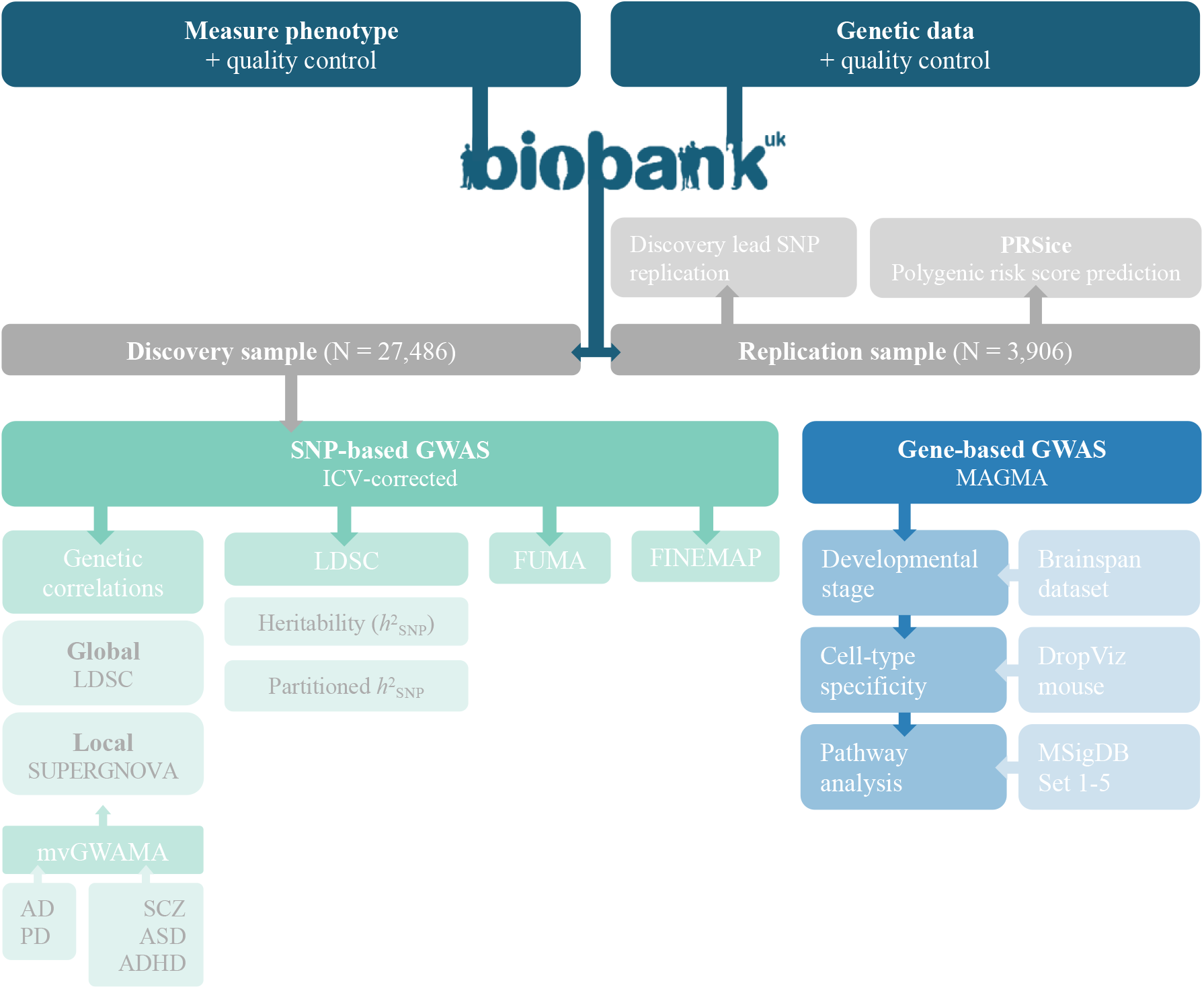
Flowchart of all data and methods used to obtain results as presented in this study. For more detailed information see Methods.

## References

1. Villanueva, R. The cerebellum and neuropsychiatric disorders. Psychiatry Res. 198, 527–532 (2012).

2. Gottwald, B., Mihajlovic, Z., Wilde, B. & Mehdorn, H. M. Does the cerebellum contribute to specific aspects of attention? Neuropsychologia 41, 1452–1460 (2003).

3. Ravizza, S. M. et al. Cerebellar damage produces selective deficits in verbal working memory. Brain 129, 306–320 (2006).

4. Gillig, P. M. & Sanders, R. D. Psychiatry, neurology, and the role of the cerebellum. Psychiatry (Edgmont). 7, 38–43 (2010).

5. Posthuma, D. et al. Multivariate genetic analysis of brain structure in an extended twin design. Behav. Genet. 30, 311–319 (2000).

6. Silventoinen, K. et al. Heritability of Adult Body Height: A Comparative Study of Twin Cohorts in Eight Countries. Twin Res. 6, 399–408 (2003).

7. Smoller, J. W. et al. Psychiatric genetics and the structure of psychopathology. Mol. Psychiatry 24, 409–420 (2019).

8. Smith, S. M. et al. An expanded set of genome-wide association studies of brain imaging phenotypes in UK Biobank. Nat. Neurosci. 24, 737–745 (2021).

9. Zhao, B. et al. Genome-wide association analysis of 19,629 individuals identifies variants influencing regional brain volumes and refines their genetic co-architecture with cognitive and mental health traits. Nat. Genet. 51, 1637–1644 (2019).

10. Chambers, T. et al. Identifying genetic variants associated with cerebellar volume in 33,265 individuals from the UK-biobank. bioRxiv 150, (2020).

11. Yang, J. et al. Genetic variance estimation with imputed variants finds negligible missing heritability for human height and body mass index. Nat. Genet. 47, 1114–1120 (2015).

12. Cross-Disorder Group of the Psychiatric Genomics Consortium. Genomic Relationships, Novel Loci, and Pleiotropic Mechanisms across Eight Psychiatric Disorders. Cell 179, 1469–1482 (2019).

13. Uffelmann, E. & Posthuma, D. Emerging methods and resources for biological interrogation of neuropsychiatric polygenic-signal. Biol. Psychiatry (2020). doi:10.1016/j.biopsych.2020.05.022

14. Choi, S. W., Mak, T. S. H. & O’Reilly, P. F. Tutorial: a guide to performing polygenic risk score analyses. Nat. Protoc. (2020). doi:10.1038/s41596-020-0353-1

15. Owen, M. J. & Williams, N. M. Explaining the missing heritability of psychiatric disorders. World Psychiatry 20, 294–295 (2021).

16. Heintzman, N. D. et al. Histone modifications at human enhancers reflect global cell-type-specific gene expression. Nature 459, 108–112 (2009).

17. Creyghton, M. P. et al. Histone H3K27ac separates active from poised enhancers and predicts developmental state. Proc. Natl. Acad. Sci. U. S. A. 107, 21931–21936 (2010).

18. Hnisz, D. et al. Super-enhancers in the control of cell identity and disease. Cell 155, 934 (2013).

19. Pott, S. & Lieb, J. D. What are super-enhancers? Nat. Genet. 47, 8–12 (2015).

20. Igolkina, A. A. et al. H3K4me3, H3K9ac, H3K27ac, H3K27me3 and H3K9me3 Histone Tags Suggest Distinct Regulatory Evolution of Open and Condensed Chromatin Landmarks. Cells 8, 1–16 (2019).

21. Frank, C. L. et al. Regulation of chromatin accessibility and Zic binding at enhancers in the developing cerebellum. Nat. Neurosci. 18, 647–656 (2015).

22. Watanabe, K., Taskesen, E., van Bochoven, A. & Posthuma, D. Functional mapping and annotation of genetic associations with FUMA. Nat. Commun. 8, 1826 (2017).

23. Roadmap Epigenomics Consortium et al. Integrative analysis of 111 reference human epigenomes. Nature 518, 317–329 (2015).

24. Watanabe, K. et al. A global overview of pleiotropy and genetic architecture in complex traits. Nat. Genet. (2019). doi:10.1101/500090

25. Boudjadi, S., Chatterjee, B., Sun, W., Vemu, P. & Barr, F. G. The expression and function of PAX3 in development and disease. Gene 666, 145–157 (2018).

26. Jara, J. et al. Pax3 induces neural circuit repair through a developmental program of directed axon outgrowth. bioRxiv (2021).

27. Kircher, M. et al. A general framework for estimating the relative pathogenicity of human genetic variants. Nat. Genet. 46, 310–315 (2014).

28. Boyle, A. P. et al. Annotation of functional variation in personal genomes using RegulomeDB. Genome Res. 22, 1790–1797 (2012).

29. van de Bunt, M., Cortes, A., Brown, M. A., Morris, A. P. & McCarthy, M. I. Evaluating the Performance of Fine-Mapping Strategies at Common Variant GWAS Loci. PLoS Genet. 11, 1–14 (2015).

30. Benner, C. et al. FINEMAP: Efficient variable selection using summary data from genome-wide association studies. Bioinformatics 32, 1493–1501 (2016).

31. Lee, J. & Cho, Y. Potential roles of stem cell marker genes in axon regeneration. Exp. Mol. Med. 53, 1–7 (2021).

32. Weiss, K. et al. Haploinsufficiency of ZNF462 is associated with craniofacial anomalies, corpus callosum dysgenesis, ptosis, and developmental delay. Eur. J. Hum. Genet. 25, 946–951 (2017).

33. Liberzon, A. et al. Molecular signatures database (MSigDB) 3.0. Bioinformatics 27, 1739–1740 (2011).

34. Saunders, A. et al. Molecular Diversity and Specializations among the Cells of the Adult Mouse Brain. Cell 174, 1015–1030.e16 (2018).

35. Miller, J. A. et al. Transcriptional landscape of the prenatal human brain. Nature 508, 199–206 (2014).

36. Phillips, J. R., Hewedi, D. H., Eissa, A. M. & Moustafa, A. A. The cerebellum and psychiatric disorders. Front. public Heal. 3, 66 (2015).

37. Wu, T. & Hallett, M. The cerebellum in Parkinson’s disease. Brain 136, 696–709 (2013).

38. Jacobs, H. I. L. et al. The cerebellum in Alzheimer’s disease: Evaluating its role in cognitive decline. Brain 141, 37–47 (2018).

39. Jansen, I. E. et al. Genome-wide meta-analysis identifies new loci and functional pathways influencing Alzheimer’s disease risk. Nat. Genet. 51, 404–413 (2019).

40. Li, L. et al. Lysine acetyltransferase 8 is involved in cerebral development and syndromic intellectual disability. J. Clin. Invest. 130, 1431–1445 (2020).

41. Nativio, R. et al. Dysregulation of the epigenetic landscape of normal aging in Alzheimer’s disease. Nat. Neurosci. 21, 1018 (2018).

42. Hackinger, S. et al. Evidence for genetic contribution to the increased risk of type 2 diabetes in schizophrenia. Transl. Psychiatry 8, (2018).

43. Ma, C., Gu, C., Huo, Y., Li, X. & Luo, X. J. The integrated landscape of causal genes and pathways in schizophrenia. Transl. Psychiatry 8, (2018).

44. Hoffmann, A. & Spengler, D. Chromatin remodeling complex NuRD in neurodevelopment and neurodevelopmental disorders. Front. Genet. 10, (2019).

45. Oxvig, C. The role of PAPP-A in the IGF system: location, location, location. J. Cell Commun. Signal. 9, 177–187 (2015).

46. Sveinsdóttir, K. et al. Impaired Cerebellar Maturation, Growth Restriction, and Circulating Insulin-Like Growth Factor 1 in Preterm Rabbit Pups. Dev. Neurosci. 39, 487–497 (2017).

47. Wrigley, S., Arafa, D. & Tropea, D. Insulin-like growth factor 1: At the crossroads of brain development and aging. Front. Cell. Neurosci. 11, 1–15 (2017).

48. Bondy, C., Werner, H., Roberts, C. T. & LeRoith, D. Cellular pattern of type-I insulin-like growth factor receptor gene expression during maturation of the rat brain: Comparison with insulin-like growth factors I and II. Neuroscience 46, 909–923 (1992).

49. Frontera, J. L. & Léna, C. When the cerebellum holds the starting gun. Neuron 109, 2207–2209 (2021).

50. Bach, M. A., Shen-Orr, Z., Lowe, W. L., Roberts, C. T. & Leroith, D. Insulin-like growth factor I mRNA levels are developmentally regulated in specific regions of the rat brain. Mol. Brain Res. 10, 43–48 (1991).

51. Hansen-Pupp, I. et al. Postnatal decrease in circulating insulin-like growth factor-I and low brain volumes in very preterm infants. J. Clin. Endocrinol. Metab. 96, 1129–1135 (2011).

52. Ye, P., Xing, Y., Dai, Z. & D’Ercole, A. J. In vivo actions of insulin-like growth factor-I (IGF-I) on cerebellum development in transgenic mice: Evidence that IGF-I increases proliferation of granule cell progenitors. Dev. Brain Res. 95, 44–54 (1996).

53. Kumar, R. et al. Purkinje cell-specific males absent on the first (mMof) gene deletion results in an ataxia-telangiectasia-like neurological phenotype and backward walking in mice. Proc. Natl. Acad. Sci. U. S. A. 108, 3636–3641 (2011).

54. Yamada, T. et al. Promoter decommissioning by the NuRD chromatin remodeling complex triggers synaptic connectivity in the mammalian brain. Neuron 83, 122–134 (2014).

55. Bipolar Disorder and Schizophrenia Working Group of the Psychiatric Genomics Consortium. Genomic Dissection of Bipolar Disorder and Schizophrenia, Including 28 Subphenotypes. Cell 173, 1705–1715.e16 (2018).

56. Visscher, P. M. et al. 10 Years of GWAS Discovery: Biology, Function, and Translation. Am. J. Hum. Genet. 101, 5–22 (2017).

57. Martin, A. R. et al. Human Demographic History Impacts Genetic Risk Prediction across Diverse Populations. Am. J. Hum. Genet. 100, 635–649 (2017).

58. Marees, A. T. et al. Genetic correlates of socio-economic status influence the pattern of shared heritability across mental health traits. Nat. Hum. Behav. (2021). doi:10.1038/s41562-021-01053-4

59. Miller, K. L. et al. Multimodal population brain imaging in the UK Biobank prospective epidemiological study. Nat. Neurosci. 19, 1523–1536 (2016).

60. Jansen, P. R. et al. Genome-wide meta-analysis of brain volume identifies genomic loci and genes shared with intelligence. Nat. Commun. 11, (2020).

61. Smith, S. M., Alfaro-almagro, F. & Miller, K. L. UK Biobank Brain Imaging Documentation. biobank.ctsu.ox.ac.uk/crystal/docs/brain_mri.pdf (2020).

62. Alfaro-Almagro, F. et al. Image processing and Quality Control for the first 10,000 brain imaging datasets from UK Biobank. Neuroimage 166, 400–424 (2018).

63. Purcell, S. et al. PLINK: A Tool Set for Whole-Genome Association and Population-Based Linkage Analyses. Am. J. Hum. Genet. 81, 559–575 (2007).

64. Abraham, G., Qiu, Y. & Inouye, M. FlashPCA2: principal component analysis of Biobank-scale genotype datasets. Bioinformatics 33, 2776–2778 (2017).

65. Alfaro-Almagro, F. et al. Confound modelling in UK Biobank brain imaging. Neuroimage 117002 (2020). doi:10.1016/j.neuroimage.2020.117002

66. Bulik-Sullivan, B. et al. LD score regression distinguishes confounding from polygenicity in genome-wide association studies. Nat. Genet. 47, 291–295 (2015).

67. Finucane, H. K. et al. Partitioning heritability by functional annotation using genome-wide association summary statistics. Nat. Genet. 47, 1228–1235 (2015).

68. Altshuler, D. L. et al. The 1000 Genomes Project Consortium: A map of human genome variation from population-scale sequencing. Nature 467, 1061–1073 (2010).

69. Ardlie, K. G. et al. The Genotype-Tissue Expression (GTEx) pilot analysis: Multitissue gene regulation in humans. Science (80-.). 348, 648–660 (2015).

70. Ramasamy, A. et al. Genetic variability in the regulation of gene expression in ten regions of the human brain. Nat. Neurosci. 17, 1418–1428 (2014).

71. Schaid, D. J., Chen, W. & Larson, N. B. From genome-wide associations to candidate causal variants by statistical fine-mapping. Nat. Rev. Genet. 19, 491–504 (2018).

72. Benner, C. et al. Prospects of Fine-Mapping Trait-Associated Genomic Regions by Using Summary Statistics from Genome-wide Association Studies. Am. J. Hum. Genet. 101, 539–551 (2017).

73. de Leeuw, C. A., Mooij, J. M., Heskes, T. & Posthuma, D. MAGMA: Generalized Gene-Set Analysis of GWAS Data. PLoS Comput. Biol. 11, 1–19 (2015).

74. Koopmans, F. et al. SynGO: An Evidence-Based, Expert-Curated Knowledge Base for the Synapse. Neuron 103, 217–234.e4 (2019).

75. Watanabe, K., Umićević Mirkov, M., de Leeuw, C. A., van den Heuvel, M. P. & Posthuma, D. Genetic mapping of cell type specificity for complex traits. Nat. Commun. 10, 3222 (2019).

76. Bulik-Sullivan, B. et al. An atlas of genetic correlations across human diseases and traits. Nat. Genet. 47, 1236–1241 (2015).

77. Grove, J. et al. Identification of common genetic risk variants for autism spectrum disorder. Nat. Genet. 51, 431–444 (2019).

78. Demontis, D. et al. Discovery of the first genome-wide significant risk loci for attention deficit/hyperactivity disorder. Nat. Genet. 51, 63–75 (2019).

79. The Schizophrenia Working Group of the Psychiatric Genomics Consortium. Mapping genomic loci prioritises genes and implicates synaptic biology in schizophrenia. medRxiv (2020).

80. Nalls, M. A. et al. Identification of novel risk loci, causal insights, and heritable risk for Parkinson’s disease: a meta-analysis of genome-wide association studies. Lancet Neurol. 18, 1091–1102 (2019).

81. Zhang, Y. et al. SUPERGNOVA: local genetic correlation analysis reveals heterogeneous etiologic sharing of complex traits. Genome Biol. 22, (2021).

82. Yengo, L., Yang, J. & Visscher, P. M. Expectation of the intercept from bivariate LD score regression in the presence of population stratification. bioRxiv 310565 (2018). doi:10.1101/310565

83. Okbay, A. et al. Genome-wide association study identifies 74 loci associated with educational attainment. Nature 533, 539–542 (2016).

84. Palmer, C. & Pe’er, I. Statistical Correction of the Winner’s Curse Explains Replication Variability in Quantitative Trait Genome-Wide Association Studies. PLoS Genet. 13, 1–18 (2017).

85. Zhong, H. & Prentice, R. L. Bias-reduced estimators and confidence intervals for odds ratios in genome-wide association studies. Biostatistics 9, 621–634 (2008).

86. Choi, S. W. & O’Reilly, P. F. PRSice-2: Polygenic Risk Score software for biobank-scale data. Gigascience 8, 1–6 (2019).

